# U-CIE [/juː ‘siː/]: Color encoding of high-dimensional data

**DOI:** 10.1101/2021.12.02.470966

**Authors:** Mikaela Koutrouli, John H. Morris, Lars J. Jensen

## Abstract

Data visualization is essential to discover patterns and anomalies in large high-dimensional datasets. New dimensionality reduction techniques have thus been developed for visualizing omics data, in particular from single-cell studies. However, jointly showing several types of data, e.g. single-cell expression and gene networks, remains a challenge. Here, we present ‘U-CIE, a visualization method that encodes arbitrary high-dimensional data as colors using a combination of dimensionality reduction and the CIELAB color space to retain the original structure to the extent possible. U-CIE first uses UMAP to reduce high-dimensional data to three dimensions, partially preserving distances between entities. Next, it embeds the resulting three-dimensional representation within the CIELAB color space. This color model was designed to be perceptually uniform, meaning that the Euclidean distance between any two points should correspond to their relative perceptual difference. Therefore, the combination of UMAP and CIELAB thus results in a color encoding that captures much of the structure of the original high-dimensional data. We illustrate its broad applicability by visualizing single-cell data on a protein network and metagenomic data on a world map and on scatter plots.

## INTRODUCTION

Today large, high-dimensional datasets are abundant in biomedicine. Data visualization is thus crucial both for discovering patterns in data and for subsequently communicating the insights. With this motivation, we present U-CIE [/juː ‘siː/] an open-source software tool that translates high-dimensional data into colors. As color space is inherently three-dimensional, U-CIE does this in two steps: a three-dimensional approximation of the high-dimensional data is produced using a dimensionality reduction method, and this approximation is next fitted into a suitable color space (Figure 1). Each high-dimensional input vector, be it the expression profile of a gene or the abundance profile of an organism, is thereby converted into a color. These colors can subsequently be used to visualize the, e.g., genes or organisms in the context of other data, such as gene networks, phylogenetic trees, or longitude and latitude.

**Figure 1.**
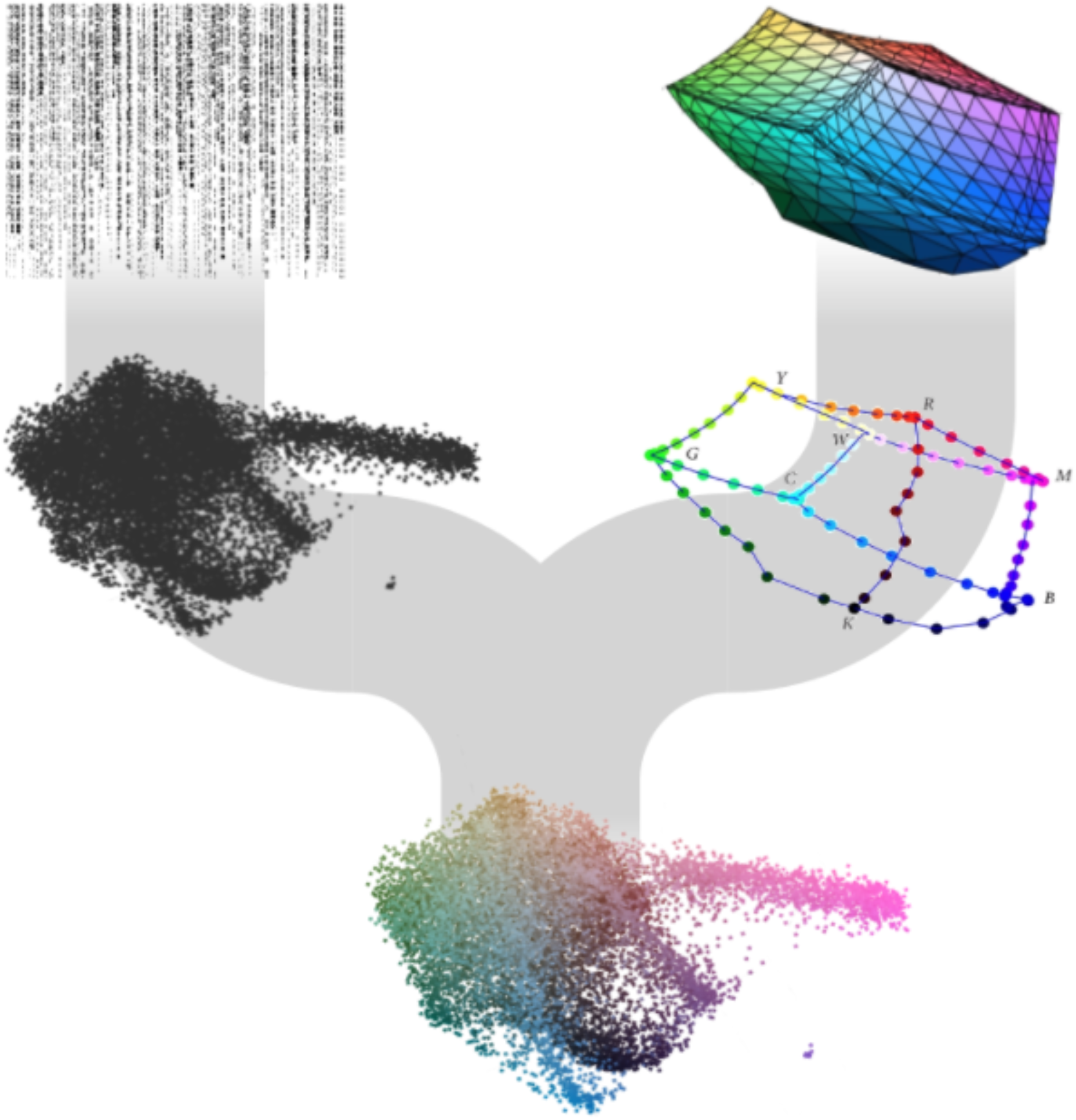
Overview of the U-CIE algorithm and application to single-cell RNAseq data (scRNA-seq). U-CIE uses a two-step process to encode high-dimensional data as colors. The first step is to use a state-of-the-art dimensionality reduction technique (UMAP) to reduce the data to three dimensions, since human color perception is inherently three-dimensional. The second step is to fit the resulting point cloud into the three-dimensional polygon that represents the displayable part of CIELAB color space. This is done as an optimization process, which shifts, rotates, and uniformly scales the point cloud to make it as large as possible while penalizing for points protruding outside the polygon.

Dimensionality reduction techniques have improved much in recent years, becoming better at preserving more of both the local and global structure in high-dimensional data by making nonlinear rather than linear transformations (1). Methods like t-SNE (2) and UMAP (3) as well as generative deep-learning models, such as variational autoencoders (4), have become particularly popular within biology, especially for visualization of single-cell data. Both t-SNE and UMAP were designed to predominantly preserve local structure by grouping neighboring data points together, which provides a very informative visualization (2,3). The main difference between t-SNE and UMAP is the interpretation of the distance between clusters. UMAP preserves pairwise Euclidean distances significantly better than t-SNE, meaning that it preserves more of the global structure along with the local. Therefore, the relative positioning of different clusters in t-SNE is not informative about the distance between them, which is why UMAP has gained popularity for visualization of single-cells studies. An alternative is to use a variational autoencoder (VAE), which is an artificial neural network that is trained to compress the data to a lower dimensional representation from which the input can be reconstructed (4). Even though VAEs can be more accurate in the low-dimensional representation, they are not commonly used yet, since they are more difficult to use than t-SNE and UMAP.

Colors are everywhere and we all have an intuitive understanding of colors; however, it is surprisingly difficult to represent them in a way that accurately reflects how similar humans perceive any two colors to be. None of the most commonly used color representations, such as RGB, approximate human vision. A notable exception is the CIELAB color model (5), which was designed to be largely perceptually uniform, meaning that the Euclidean distance between any two points should correspond to their relative perceptual difference. This makes it particularly useful for our application, since we want to visualize positions as colors, similar to what has previously been tried for self-organizing maps (6). CIELAB represents colors using three values: perceptual lightness (L*) and two pairs of complementary colors, namely green–red (a*) and blue–yellow (b*). To take advantage of the properties of CIELAB, we need to convert the three-dimensional coordinates obtained from dimensionality reduction into L*a*b* coordinates. This is nontrivial, because not all combinations of L*, a*, and b* result in colors that can be displayed on a computer monitor.

Here, we present the U-CIE visualization method for encoding high-dimensional data as colors. We first use UMAP to turn the data into a three-dimensional point cloud, as UMAP partially preserves global Euclidean distances. Next, we use an optimization algorithm to fit the point cloud within the displayable part of CIELAB. The result is an encoding of the input data, where each data point has been assigned a color, and similarity between colors reflect the distance between the original points. We illustrate the broad applicability of U-CIE by using it to visualize single-cell expression data on a protein network and microbiome composition of ocean water samples on a world map. U-CIE is freely available both as a web resource (https://u-cie.jensenlab.org) and as an R package.

## MATERIAL AND METHODS

### Algorithm overview

U-CIE color encodes high-dimensional data using CIELAB color space in two stages: i) dimensionality reduction (can be skipped if data are already three-dimensional) and ii) fitting the resulting point cloud inside the displayable part of CIELAB (Figure 1). The latter is a precomputed polygon, constructed by converting the RGB cube to CIELAB coordinates.

### Dimensionality reduction

Unless the input data is already three-dimensional, U-CIE will start by reducing the data to three dimensions. There are three different tracks for doing so: ‘Single cells’, ‘High dimensional data’, and ‘Distance matrix’. The ‘Single cells’ track follows the Seurat (7) pipeline to reduce dimensionality and produce 3D UMAP coordinates for the genes of the dataset. To do so, we first transpose the input dataset to have genes as columns and cells as rows and create a Seurat Object from the counts. Without first scaling or centering the data, we use Seurat to log2 transform our data using the (option “LogNormalize”), run PCA with 50 dimensions, calculate 50 neighbors for each gene, and find clusters with the Louvain algorithm. Finally, we apply UMAP to end up with three dimensions. The ‘High dimensional’ track uses the uwot library to again run PCA with 50 dimensions and apply UMAP. Finally, the ‘Distance matrix’ track takes a square matrix and uses Python umap-learn package.

### Fitting the point cloud inside the CIELAB RGB polygon

To be able to handle large datasets with many points, we first use the R ‘chull’ function to construct the convex hull of the point cloud. This dramatically speeds up the subsequent optimization step, typically by several orders of magnitude, as it only has to consider the few points on the convex hull rather than all points in the input dataset. Next, we use the Nelder–Mead simplex optimization algorithm (8) to fit the convex hull of the point cloud inside the CIELAB RGB polygon. The algorithm is allowed to shift along and rotate around three axes (L*, a*, and b*) and uniformly scale the point cloud convex hull. The objective function to be optimized is the size of the point cloud minus a penalty term for points falling outside the polygon. In other words, it aims to make the point cloud as large as possible while still fitting within the color space. To avoid local optima, we run the algorithm with 25 different sets of initial rotations. The user can optionally provide different weights for the axes in the objective, thus prioritizing spreading out the points along certain axes.

### User Interface

The web interface was constructed using R/Shiny and JavaScript. An interactive guided tutorial is available through the web interface as well as a YouTube video explaining the idea of U-CIE.

After uploading and processing data via one of the tracks described above, the data can be visualized in two ways. The ‘3D view’ shows the main result of following the U-CIE pipeline, namely the 3D cloud of points colored according to the best solution found by the optimization algorithm. The view also has a table of alternative color solutions from the optimizations, which can be selected and displayed instead. The ‘2D projections’ tab provides two interactive 2D plots, showing the same 3D point cloud projected onto the L*a* and L*b* axes, respectively. There users can see how the points are spread within the polygon. The view also has a table, which allows the user to select regions within the plots to identify the data points or search the data points by name to locate them within the cloud. Finally, the ‘Download’ tab shows a table with both RGB hex codes and CIELAB coordinates, which can be downloaded to use the colors for further data visualization, e.g. in Cytoscape [8]. The plots and tables are made using the Plotly (9) and DataTables CRAN.R-project.org/package=data.table libraries, respectively.

### CRAN package ‘ucie’

Our algorithm is also available as a CRAN package in R. Users can download it from R with the command install.packages(“ucie”) or from the packages and the command devtools::install_github(“mikelkou/ucie”). The package returns a data frame with the names of the input data points and RGB hex colors or CIELAB coordinates. The package contains only one function with 6 arguments, 5 of which are optional. Users must provide a 3-column dataset that will be encoded as colors. The other optional arguments allow users to alter the axis weights, a final scaling factor, and whether they want RGB hex codes or CIELAB coordinates. The output is a data frame with the names of the input data points and the corresponding colors.

## RESULTS AND DISCUSSION

### U-CIE [/juː ‘siː/] an open-source software tool

To allow anyone to easily use U-CIE, we have made it available as an interactive web resource (https://u-cie.jensenlab.org) along with a guided tutorial. The web interface offers four different tracks: ‘Single cells’, ‘High dimensional’, ‘Distance matrix’, and ‘3D data’. All use UMAP to first reduce dimensionality, except from the last track, which allows the user to upload data that is already three-dimensional, possibly produced using another dimensionality reduction method. In Supplementary Figure S1 we show how an analysis can be performed using the web interface. First, we upload the matrix with expression counts to the ‘Single cells’ track, which assigns colors to genes. A preview of the uploaded data is first shown, allowing us to verify that it was parsed correctly before starting the analysis. As soon as the analysis has finished, we gain access to interactive visualization panels that show the color encoding in 3D and as 2D projections, controls to alter the color encoding, and the option to download a tab-delimited file with the gene names and their colors.

U-CIE is also available as an R package. Since users of this package will be working in the R environment, we give them full flexibility to use any dimensionality reduction method for creating the three-dimensional representation of the input data. The R package thus takes this as input and performs the optimization to fit the data into CIELAB coordinates. The output is a data frame with the names of the input data and the hex codes of the colors or the CIELAB coordinates. This data frame can be directly used with R or saved as a file locally for use in other visualization tools. Both the R package and the web resource are available under the open source MIT License.

### scRNA-seq data on physical protein complexes

Having a color encoding of high-dimensional data is useful whenever you want to visualize the data in the context of some other information, since it frees up the spatial coordinates. For example, transcriptomics and proteomics data are commonly visualized onto gene/protein networks. We exemplify this use of the U-CIE method by applying it to network visualization of a scRNA-seq data of 19,097 genes across 1018 cells. We used U-CIE to convert this data to a color for each gene and mapped them on a physical protein interaction network from STRING v11.5 (10,11) (confidence cutoff 0.95) using Cytoscape (12). This allows us to graphically summarize the transcriptional regulation of the individual subunits within protein complexes. Figure 2 shows the six largest connected components of the network. Several clusters corresponding to large protein complexes with many green subunits stand out in the network, including the cytosolic and mitochondrial ribosomes, the proteasome, and the electron transfer chain complexes. As these carry out housekeeping functions it makes sense that they have very similar expression patterns across cells and thus have the same color. The electron transfer chain complexes also have a few blue subunits, which are the ones encoded by the mitochondrial chromosome. The network also contains a cluster of tan proteins, which are involved in core cell-cycle processes, such as the cyclin-dependent protein kinases and DNA replication complexes. This example illustrates that U-CIE is able to assign colors to genes based on scRNA-seq data in a biologically meaningful way.

**Figure 2.**
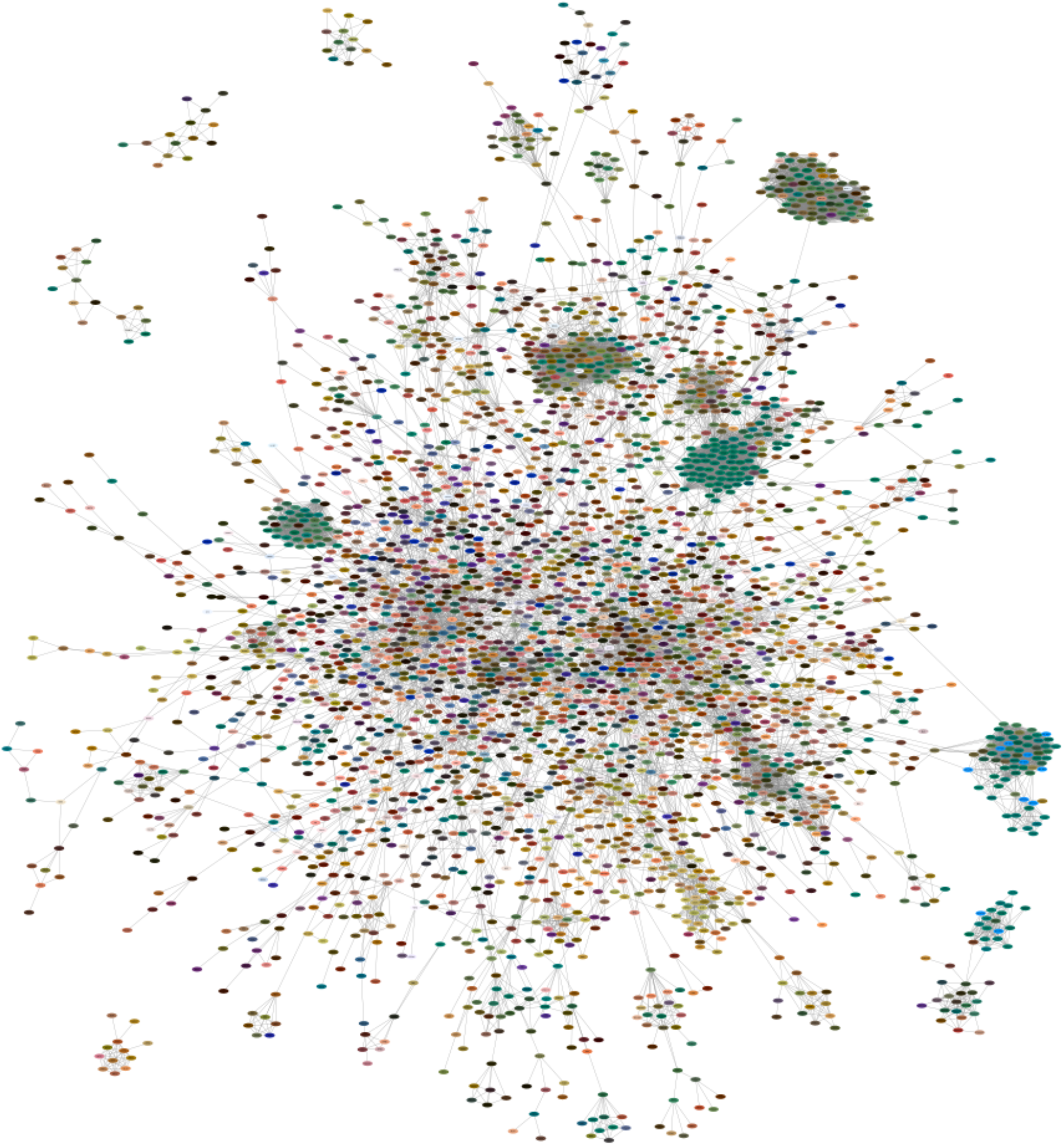
Color encoding the scRNA-seq on physical protein complexes. To illustrate how U-CIE can be used to visualize expression data on protein networks, we downloaded a published scRNA-seq dataset with a read-count matrix of 1018 cells (13). We applied U-CIE to the transposed matrix (genes as columns, cells as rows), exported the colors, and visualized them on a physical protein–protein interaction network from the STRING database (7) (confidence cutoff 0.95) using Cytoscape (8). In the figure, we show the six largest connected components. The network contains several large dark green clusters; these correspond to large complexes of housekeeping proteins, such as the cytosolic and mitochondrial ribosomes, the proteasome, and the electron transport chain complexes. The latter also contain some blue subunits, which are the proteins encoded by the mitochondrial genome. The less obvious cluster of tan nodes corresponds to core cell-cycle proteins, including cyclin-dependent protein kinases and the DNA replication complexes.

### Microbiome compositions on a world map

U-CIE is not limited to single-cell data. Figure 3 shows a completely different use case, namely visualization of microbiome compositions. Specifically, we used U-CIE to color the sampling stations of the Tara Oceans project (14) based on their 16S taxonomic profiles from surface water samples. Combining these colors with the longitudes and latitudes of the sampling stations enables us to show the microbiome compositions on a world map (Figure 3A). This makes it immediately clear that the samples from some oceans have quite similar microbiome composition. For example, samples from the Mediterranean Sea tend to be blue while samples from the Indian Ocean tend to be yellow (Figure 3A).

**Figure 3.**
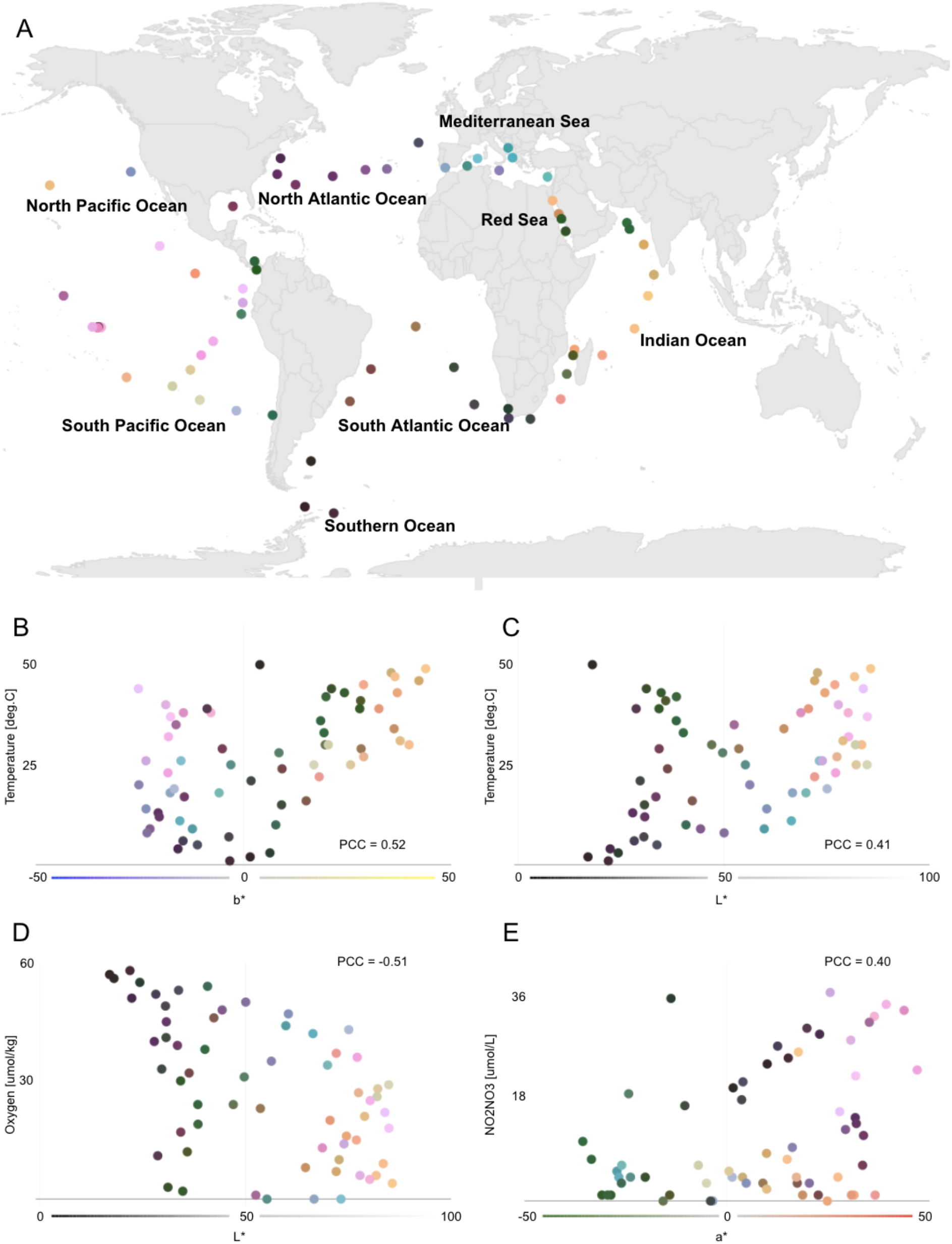
Color encoding the microbiome compositions of ocean surface water samples. 16S rRNA microbiome composition of surface water samples from the Tara Oceans (14) sampling stations were encoded as colors using U-CIE. The resulting colors were mapped onto the geographic locations of the sampling stations (panel A). The blue–yellow (b*) axis correlates with water temperature (panel B), the lightness (L*) correlates positively with temperature dissolved (panel C) and negatively with oxygen concentration (panel D), and the green–red axis (a*) correlates with dissolved nitrite/nitrate concentration (panel E).

The colors produced by U-CIE can also be interpreted in terms of environmental factors that drive microbiome composition. We did this by correlating each factor measured in the Tara Oceans project with the individual CIELAB color axes. The strongest correlation found is that the blue–yellow axis (b*) correlates with water temperature (PCC=0.52; Figure 3B). This agrees with the observation in the original publication that temperature correlates strongly with microbiome composition (14). The lightness (L*) also correlates with water temperature (PCC=0.41; Figure 3C) but shows a stronger negative correlation with the dissolved oxygen concentration (PCC=-0.51; Figure 3D). Together, these axes thus correctly capture, from 16S abundance profiles, that the Indian ocean is warm with low oxygen, the North Atlantic Ocean is cold with high oxygen, and the Pacific Ocean has low oxygen and highly varying temperatures. Finally, the green–red axis (a*) correlates with dissolved nitrite/nitrate concentration (panel E); for example, the middle of the Pacific Ocean is a high-nitrogen environment, whereas the waters around Panama have low nitrogen.

## SUMMARY

We have developed a new visualization tool, U-CIE, which allows arbitrary high-dimensional data to be encoded as colors. The method first uses existing dimensionality reduction techniques, e.g. UMAP, to reduce the input data to three dimensions, and next embeds this representation of the data within the CIELAB color space. We illustrate the usefulness of U-CIE by applying it to i) visualization of scRNA-seq data on a physical protein interaction network and ii) visualization of microbiome composition from sampling stations on a world map. U-CIE is available both as a web resource at https://u-cie.jensenlab.org/ and as an R package.

## DATA AVAILABILITY

Each track offers a different small example dataset. The ‘Single cells’ track uses a single-cell RNA-seq dataset of human embryonic stem cells (GEO accession code GSE75748) (13). The ‘High dimensional’ track uses an Affymetrix time course dataset of *Drosophila melanogaster* embryogenesis (15). The ‘Distance matrix’ example uses the bacterial species tree available from the iTOL web resource (https://itol.embl.de) (16). For the ‘3D Data’ track we used the same dataset as in the ‘High dimensional’ track, reduced to three dimensions with locally linear embedding. Finally, the dataset used to create Figure 2 comes from (14) and can be downloaded from http://ocean-microbiome.embl.de.

## CODE AVAILABILITY

The U-CIE source code is available at https://github.com/mikelkou/U-CIE_Web_Resource. The R package ‘ucie’ code is available at https://github.com/mikelkou/ucie, through CRAN (install.packages(“ucie”)), and through devtools (devtools install_github(“mikelkou/ucie”))

## ACKNOWLEDGEMENT

The authors acknowledge Katerina Nastou for input on figures and Casper H. Blaauw for input on the R code.

## REFERENCES

1. LJ van der Maaten, PostmaP E, J van den Herik. Dimensionality reduction: a comparative review. Journal of Machine Learning Research. 2009;10.1(41):66–71.

2. van der Maaten, LJ, Hinton, G. MG, Geoffrey. Visualizing Data using t-SNE. J Mach Learn Res. 2008;(9):2579––2605.

3. McInnes L, Healy J, Melville J. UMAP: Uniform Manifold Approximation and Projection for Dimension Reduction. ArXiv180203426 Cs Stat [Internet]. 2020 Sep 17 [cited 2021 Apr 8]; Available from: http://arxiv.org/abs/1802.03426

4. Hinton GE, Salakhutdinov RR. Reducing the Dimensionality of Data with Neural Networks. Science. 2006 Jul 28;313(5786):504–7.

5. Internationale Beleuchtungskommission, editor. Colorimetry. 3rd ed. Wien: Comm. Internat. de l’eclairage; 2004. 72 p. (Publication / CIE).

6. Wang FY, Takatsuka M. Self-organizing Map (SOM) Based Data Navigation for Identifying Shape Similarities of Graphic Logos. Neural Process Lett. 2015 Jun;41(3):325–39.

7. Hao Y, Hao S, Andersen-Nissen E, Mauck WM, Zheng S, Butler A, et al. Integrated analysis of multimodal single-cell data [Internet]. Genomics; 2020 Oct [cited 2021 Mar 3]. Available from: http://biorxiv.org/lookup/doi/10.1101/2020.10.12.335331

8. John A. Nelder and Roger Mead. A simplex method for function minimization. Computer Journal. 1965;7:308––313.

9. Sievert C. Interactive web-based data visualization with R, plotly, and shiny. Boca Raton, FL: CRC Press, Taylor and Francis Group; 2020.

10. Szklarczyk D, Gable AL, Nastou KC, Lyon D, Kirsch R, Pyysalo S, et al. The STRING database in 2021: customizable protein–protein networks, and functional characterization of user-uploaded gene/measurement sets. Nucleic Acids Res. 2021 Jan 8;49(D1):D605–12.

11. Doncheva NT, Morris JH, Gorodkin J, Jensen LJ. Cytoscape StringApp: Network Analysis and Visualization of Proteomics Data. J Proteome Res. 2019 Feb 1;18(2):623–32.

12. Shannon P. Cytoscape: A Software Environment for Integrated Models of Biomolecular Interaction Networks. Genome Res. 2003 Nov 1;13(11):2498–504.

13. Chu L-F, Leng N, Zhang J, Hou Z, Mamott D, Vereide DT, et al. Single-cell RNA-seq reveals novel regulators of human embryonic stem cell differentiation to definitive endoderm. Genome Biol. 2016 Dec;17(1):173.

14. Sunagawa S, Coelho LP, Chaffron S, Kultima JR, Labadie K, Salazar G, et al. Structure and function of the global ocean microbiome. Science. 2015 May 22;348(6237):1261359.

15. Tomancak P, Beaton A, Weiszmann R, Kwan E, Shu S, Lewis SE, et al. Systematic determination of patterns of gene expression during Drosophila embryogenesis. Genome Biol. 2002;3(12):research0088.1.

16. Letunic I, Bork P. Interactive Tree Of Life (iTOL): an online tool for phylogenetic tree display and annotation. Bioinformatics. 2007 Jan 1;23(1):127–8.

