## Supplementary Figure S1 for "U-CIE [/juː ‘siː/]: Color encoding of high-dimensional data"

### SUPPLEMENTARY DATA

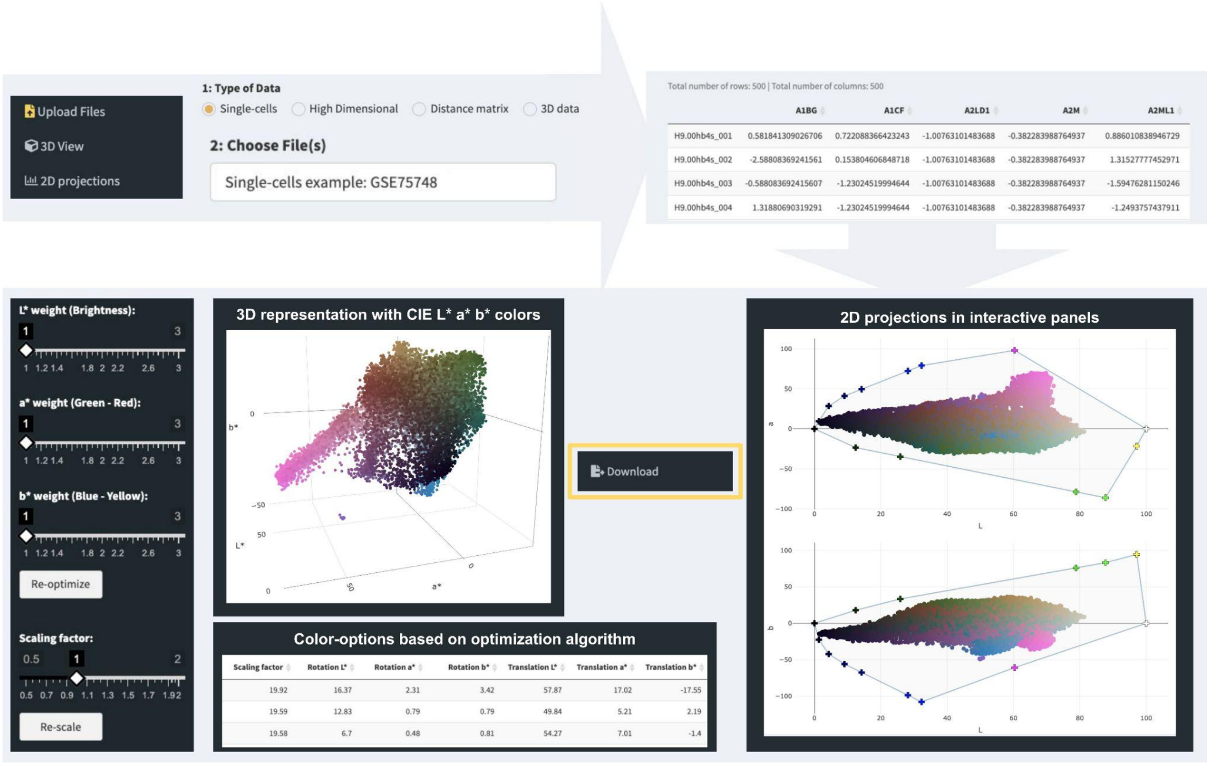


Figure S1. **Web-resource interface of U-CIE.** The web interface was constructed using R/Shiny and JavaScript. The first choice users need to make is which track is more suitable for their data. After uploading and processing data via one of the tracks, there are two outputs of the data. The '3D view' shows the best solution found by the optimization algorithm following the U-CIE pipeline, namely the 3D representation of the cloud of points colored. Alternative colors can be selected and displayed instead from the table of alternative color solutions from the optimizations. The '2D projections' tab provides two interactive 2D plots, showing the same 3D point cloud projected onto the L*a* and L*b* axes, respectively. There users can see how the points are spread within the polygon. Finally, from the 'Download' tab users can download the colors and use them for further data visualization, e.g. in Cytoscape. The plots and tables are made using the Plotly and DataTables libraries, respectively.
